# Assessing bias and robustness of social network metrics using GPS based radio-telemetry data

**DOI:** 10.1101/2023.03.30.534779

**Authors:** Prabhleen Kaur, Simone Ciuti, Federico Ossi, Francesca Cagnacci, Nicolas Morellet, Anne Loison, Kamal Atmeh, Philip McLoughlin, Adele K. Reinking, Jeffrey L. Beck, Anna C. Ortega, Matthew Kauffman, Mark S. Boyce, Michael Salter-Townshend

## Abstract

1. Social network analysis of animal societies allows scientists to test hypotheses about social evolution, behaviour, dynamical processes, and transmission events such as the spread of disease. However, the accuracy of estimated social network metrics depends on the proportion of individuals sampled, actual sample size, and frequency of observations. Robustness of network metrics derived from a sample has thus far been examined through various simulation studies. However, simulated data do not necessarily reflect the nuances of real empirical data.
2. We used some of the largest available GPS telemetry relocation datasets from five species of ungulates characterised by different behavioural and ecological traits and living in distinct environmental contexts to study the bias and robustness of social network metrics. We introduced novel statistical methods to quantify the uncertainty in network metrics obtained from a partial population suited to autocorrelated data such as telemetry relocations. We analysed how social network metrics respond to down-sampling from the observed data and applied pre-network data permutation techniques, a bootstrapping approach, correlation, and regression analyses to assess the stability of network metrics when based on samples of a population.
3. We found that global network metrics like density remain robust when the sample size is lowered, whereas some local network metrics, such as eigenvector centrality, are entirely unreliable when a large proportion of the population is not monitored. We show how to construct confidence intervals around the point estimates of these metrics representing the uncertainty as a function of the number of nodes in the network.
4. Our uncertainty estimates enable the statistical comparison of social network metrics under different conditions, such as analysing daily and seasonal changes in the density of a network. Despite the striking differences in the ecology and sociality among the five different ungulate species, the various social network metrics behave similarly under downsampling, suggesting that our approach can be applied to a wider range of species across vertebrates. Our methods can guide methodological decisions about animal social network research (e.g., sampling design and sample sizes) and allow more accurate ecological inferences from the available data.

## Introduction

Social network analysis (SNA) has proved to be a valuable toolkit for biologists to understand diverse interactions among animal communities and their effect on the environment (***James et al., 2009***; ***Sosa et al., 2021a***; ***Kulahci et al., 2016***). SNA also helps in understanding how environmental factors influence the structure of animal communities (***Hock and Fefferman, 2011***; ***Krause et al., 2016***, ***2010***; ***Albery et al., 2021***) and informs how minor changes at the individual level propagate changes in the overall behaviour of the population (***Aplin et al., 2012***; ***Farine et al., 2015***; ***Shimada and Sueur, 2014***) which in turn, contributes to informing epidemiological models, implementing customized measures for disease control, designing wildlife conservation policies, and resource allocation (***Egan et al., 2023***; ***Silk et al., 2017***). The term SNA is used to refer to the analysis of network data, for which there is an expanding set of statistical models and inferential procedures (see ***Salter-Townshend et al. (2012)*** for an introduction). Under this definition, the observed interactions between individuals are taken as fact and the goal is to summarise the data and make various inferences from it about the structure of the population of individuals and perhaps also of the behaviour of each individual. However, the term SNA may more broadly refer to the algorithmic construction of interactions data from direct observations of individuals, followed by the application of these bespoke SNA based methods. It is therefore important to clarify since the beginning that in our paper we demonstrate some key considerations that should be made in both the construction and analysis of animal social networks (***Croft et al., 2008***; ***Whitehead, 2008***; ***Krause et al., 2007***; ***Lusseau et al., 2008***) with a focus on assessing the reliability and robustness of the more commonly reported network metrics, as calculated on interactions data constructed from observations of individuals.

One of the fundamental requirements for performing social network analysis on animals is that a substantial portion of individuals in the population is uniquely identified and observed for a sufficient period (***Farine and Whitehead, 2015***). Recent advances in Global positioning system (GPS) telemetry technology have led to a significant boost in animal tracking and enabled wildlife ecologists to monitor and map minute details of animal movements, including those of highly cryptic species (***Smith and Pinter-Wollman, 2021***; ***Cagnacci et al., 2010***; ***Crofoot, 2021***; ***Webber and Wal, 2019***). However, deploying GPS devices can be expensive. Commercial wildlife devices cost thousands of dollars each, depending on the study species and required features. Even the recent low-priced solutions cost more than $100 per collar (***Foley and Sillero-Zubiri, 2020***), which restricts the number of individuals monitored simultaneously. In addition to this, issues in marking all individuals of a population include constraints on the human effort required (which is also costly), geographical constraints of some or all individuals in the population where capture methods do not work, personality traits of individuals as some individuals are capture-shy (***Biro and Dingemanse, 2009***), and ethical and other issues raised by some stakeholder.

Therefore, data representing a large sample size is a significant limitation, especially while analysing social networks (***He et al., 2022***). This is a concern as the relations among the members obtained from a sample of GPS-tracked individuals under represent their complete set of relationships (***Croft et al., 2008***). Furthermore, missing individuals from the sample may strongly influence the sampled individuals’ social measures. Thus, relational data could be expected to respond more unreliably to sampling from a population than other data types (***Silk et al., 2015***). The relational nature of network data also causes it to violate the assumptions of independence that underlies most of the parametric statistical tests (***Farine and Whitehead, 2015***) and creates an additional challenge in using a sample of individuals to make inferences about the population.

It is therefore crucial that animal social network studies consider the robustness of current methodological approaches to data rarefaction and randomization, as both the collected data and analytical methods are prone to biases inflicted by specifics of sampling protocols or the species under study (***Sosa et al., 2021a***). Networks constructed using a random subset of population are termed as partial networks (***Silk et al., 2015***). The effect of using partial networks on the properties of individual metrics in animal social network analysis has received so far little attention (***Croft et al., 2008***; ***Perkins et al., 2009***; ***Cross et al., 2012***; ***Silk et al., 2015***) and to date, there are no established methods for estimating if a partial network is a good representation of the real social structure (***Farine and Strandburg-Peshkin, 2015***), and the associated level of uncertainty (***Bonnell and Vilette, 2021***). Previous research has been primarily focused on the impact of missing nodes on social network structure, suggesting that the network statistics derived from partial data are biased estimators of the overall network topology (***Bliss et al., 2014***). Thus, understanding how network statistics scale with sampling regime is important. Some preliminary work has been conducted to determine scaling methods, predict true network statistics from a partial knowledge of nodes, links or weights of a network, and eventually validate the results on simulated networks and twitter reply networks (***Bliss et al., 2014***)). Another simulation study (***Silk et al., 2015***) has further highlighted the importance of understanding consequences of missing random nodes from a complete animal social network. Specifically, the networks have been simulated following a typical fluid fission fusion social system to determine the precision and accuracy of measures of individual social positions based on incomplete knowledge. On the contrary of what expected, ***Silk et al. (2015)*** found that in social networks based on fluid social interactions precise inferences about individual social position can be derived even when not all individuals in a population are identifiable. Despite the importance of such findings, since then research has not progressed in this field. In particular, there is the necessity to understand the level of confidence, and associated bias, with whom the current methods adopted to estimate partial network from subsampled populations actually catch the structure of the real-world animal social network (***Farine and Whitehead, 2015***; ***Sosa et al., 2021b***; ***Silk et al., 2015***).

Our paper aims to present methods that can assess the sufficiency (i.e., based on estimated bias and uncertainty) of the available data sample to perform social network analysis and obtain a measure of accuracy for global and node-level network metrics (***Farine and Whitehead, 2015***). Our approach is particularly suited (but not limited) to telemetry relocations considering their autocorrelated structure (***Boyce et al., 2010***). For this, we present a four-step paradigm applied to GPS telemetry observations of five species of ungulates with very different ecology and living in heterogeneous ecosystems.

1. The first step is to determine if the network structure obtained from the available sample of GPS observations captures any non-random aspects of the association. For this, we generate null networks by permuting a pre-network data stream. If a specific network metric does not meet this requirement, it should be discarded by researchers in their specific study case.
2. The second step is to assess how the bias in the network summary statistics varies with a decrease in the proportion of individuals sampled. Sub-sampling from the observed network helps estimate the extent of uncertainty in the network summary statistics and provides an idea of the robustness of the available sample.
3. The third step is to explore how different the network properties would have been if the researchers had tagged a completely different set of individuals from the population. This is achieved by applying a bootstrapping technique on the subsamples of the observed network. We also assess uncertainty by obtaining confidence intervals around the values of observed network statistics with the help of bootstrapping, which is also critical when it comes to comparing networks (e.g., daily or seasonal changes in sociality, or between two populations of the same species.)
4. The fourth and final step is to check how the node-level network metrics are affected by the proportion of individuals present in the sample. We use correlation and regression analyses to assess the robustness of node-level characteristics.

We conclude our paper by outlining the methods described above and provide a step-wise protocol for ecologists on the application of these on their datasets. We have recently published a companion R software package aniSNA (***Kaur, 2023***), which serves as a ready-made toolkit for ecologists to apply the methods described in this paper in their animal social network studies.

## Materials and Methods

### Data

We collated high-frequency GPS telemetry relocations’ datasets from five species of ungulates, namely caribou (*Rangifer tarandus*), elk (*Cervus canadensis*), mule deer (*Odocoileus hemionus*), pronghorn (*Antilocapra americana*), and roe deer (*Capreolus capreolus*) belonging to four different geographical regions (Table 1). These large datasets consist of observations from a proportion of individuals sampled from the population and contain a unique animal identity number, date, time, and spatial coordinates of the observations.

**Table 1.** Summary of the data available for five species of ungulates monitored using satellite telemetry in North America and Europe.

| Species Name | Area of Observation | Centroid (Lat, Lon) | Number of Animals Observed | Total number of Observations | Duration of observation | Fix Rate |
| --- | --- | --- | --- | --- | --- | --- |
| Caribou | Saskatchewan, Canada | (57.14489, -104.3752) | 94 (F:94, M:0) | 304,607 | 03/2014 to 03/2018 | Every 5 hours |
| Elk | Rocky Mountains, Alberta, Canada | ( 49.52496 , -114.3014 ) | 171 (F:111, M:60) | 856,241 | 01/2007 to 03/2013 | Every 2 hours |
| Mule Deer | Red Desert, Wyoming, US | ( 42.24222 , -109.2664 ) | 263 (F:256, M:7) | 1,458,043 | 03/2014 to 06/2021 | Every 1-2 hours |
| Pronghorn | Red Desert, Wyoming, US | ( 41.60466 , -107.9531 ) | 159 (F:159, M:0) | 896,401 | 11/2013 to 10/2016 | Every 2 hours |
| Roe Deer | Aurignac, France | ( 43.28552 , 0.8809104 ) | 147 (F:81, M:66) | 419,165 | 01/2005 to 12/2012 | Every 10 minutes |

### Associations and Network Construction

#### Identifying Associations from Raw GPS data

We obtained network structure from the raw data stream by identifying associations between each pair. We considered a pair of individuals in the sample to be associating if the two animals were observed within *s* metres from each other and within a time frame of *t* minutes. The value of spatial threshold *s* can be chosen by applying a statistical approach to the observed data. ***He et al. (2022)*** suggest one such approach could be to use the first mode from the distribution of inter-individual distances as it likely represents socially associating individuals. The temporal threshold *t* is dictated by the fix rates in telemetry data. For example, GPS collars on animals send signals consisting of spatial coordinates after a predetermined time interval. These signals can be received a few seconds (up to a few minutes) before or after the expected time. Therefore, temporal thresholds should be chosen in such a way that it accounts for this flexibility. Researchers should generally pick a threshold based on their device accuracy, species ecology, and research question.

#### Association Index and Network Formation

An association index was calculated through a modified version of Simple Ratio Index (***Farine and Whitehead, 2015***) for GPS telemetry observations. ***He et al. (2022)*** argue that for GPS data, an observation of individual A without individual B is only informative if B is observed elsewhere simultaneously. Therefore, the denominator of the original formula should only include observations where GPS data are simultaneously available for both individuals. The modified index used in the analysis is as follows:

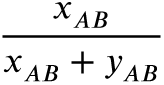

where,

*x_AB_*− No. of times when A and B are observed associating

*y_AB_*−No. of times A and B are observed within the temporal threshold but not associating

The value of the index ranged between 0 and 1, where 0 indicated that the two animals were never observed together and 1 indicated that they were always observed together. The individuals sampled from the population form the network nodes, and an association between pairs account for the edges in the networks. Each edge in the network had a weight attribute that signified the association strength calculated by the modified Simple Ratio Index described above. In this way, we obtained the network structures corresponding to each species from their raw GPS observations which also represent the complete set of relationships among the individuals tagged for each species.

### Analysis

To assess the properties of the networks, we used standard metrics common in animal social network analysis (Table 2). The local network summary statistics which provided individual level information included degree (***Shimada and Sueur, 2014***), strength (***Pike et al., 2008***), betweenness centrality (***Kanngiesser et al., 2011***; ***Aplin et al., 2012***), eigenvector centrality (***Kulahci et al., 2016***; ***Aplin et al., 2012***) and local clustering coefficient (***Pike et al., 2008***; ***Shimada and Sueur, 2014***). Density (***Ozella et al., 2021***), transitivity, and diameter are the global network metrics and provide a summary of the overall network and behaviour of the individuals as a whole. We also calculated the mean of each node’s degree (mean degree (***Shimada and Sueur, 2014***)) and strength (mean strength) and used those as global network properties.

**Table 2.**
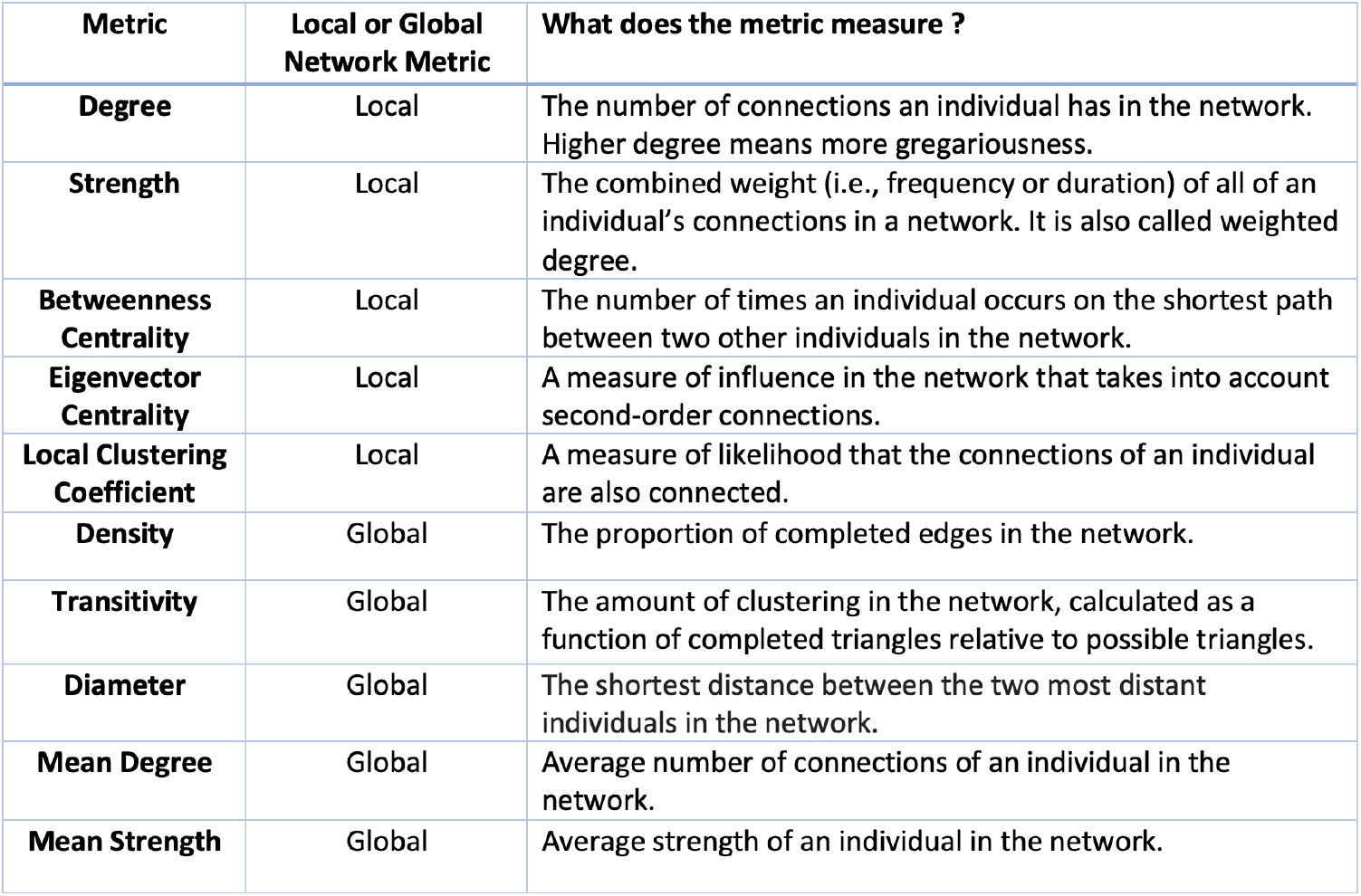
Network metrics used in the analyses

### 1. Pre-network data permutations

To assess if the interactions captured by the observed sample were genuinely caused by social preferences, we generated null models. Null models were constructed to account for non-social factors that lead to the co-occurrence of animals. In animal social network analysis, null models are broadly classified in two ways: network permutations and pre-network permutations (***Farine, 2017***). Network permutations are performed after the network is generated from the data, whereas pre-network permutations are performed on the data stream before generating networks from it. GPS telemetry observations generate data in the form of autocorrelated streams. In the permuted versions of the data, we wanted to maintain this autocorrelation structure of each individual’s movements but randomize the contacts. Therefore, we obtained pre-network datastream permutations as suggested by ***Spiegel et al. (2016)***, and Farine (2017). For each individual in the study, the tracks followed by them on each day were segmented. Then the dates on which those tracks were followed were shuffled for each individual. This methodology of permuting the pre-network data stream ensured unaffected home ranges of animals in the permuted data, but whom they came in contact with was now randomized in the null model. This also preserved the autocorrelated structure of individual tracks to ensure realistic animal movements.

For each species, we obtained 1000 permuted versions of the raw data stream, giving rise to 1000 network structures. Then, we calculated global network summary statistics for each of those networks and obtained a null distribution of values. We then compared the observed network properties to the distribution of null values, which helped determine the metrics that capture non-random aspects of the observed network.

### 2. Analysing sub-samples of the observed network

We randomly sub-sampled *m* nodes from the observed network of *N* nodes where *m* < *N* without replacement. All the associations among the sampled nodes were preserved, and the rest were dropped. This resulted in a network structure that would have been obtained if originally just these *m* individuals had been tagged from the population. In this way, we drew 100 samples of size *m* where the value of *m* ranged from 10% to 90% of the total nodes forming each network for five species. We recorded the values of global network metrics of density, mean strength, transitivity, and diameter and obtained a distribution of the values. We assessed the bias in the values of network metrics obtained from this sub-network compared to the original network. Performing this procedure across five species and for different values of *m* revealed the robust network metrics that should be adopted for social network studies on the available samples.

We also applied a sub-sampling approach on the permuted networks to determine under what sampling level the observed networks start to resemble the random networks. We sub-sampled nodes from 1000 permuted network versions without replacement at different levels ranging from 10% to 90%. Four global network metrics were calculated for each permuted version, and each level and their distribution were plotted along with the distribution of sub-sampled versions of the original network. This visualisation provided an estimate of the minimum amount of subsampling required to ensure that the network differed significantly from a random network for that species and the environment.

### 3. Bootstrapped confidence intervals

Assuming a researcher has chosen a set of network metrics appropriately, it is prudent to consider not only the point estimate derived from their data but also the uncertainty associated with it. In order to create confidence intervals to facilitate the comparison of different networks (e.g., differing sampled individuals from the same population or the same individuals’ networks computed at different times), we adapted the Bootstrap algorithms of ***Snijders and Borgatti (1999)***. Similar to the algorithms used in SOCPROG (***Whitehead, 2009***) and UCINET (***Borgatti et al., 2014***), this algorithm sampled nodes in the network with replacement for each of B=1000 Bootstrap replications. In each replication, edges between two different resampled nodes were retained, whereas edges between the same node resampled twice were sampled uniformly at random from the set of all original edges. Therefore, each bootstrap replication network comprised the same number of nodes (animals) as the original network; however, some of the original nodes were absent, some were present once, and some more than once.

Bootstrapping has been used to infer uncertainty in animal social networks (see ***Lusseau et al. (2008)***, ***Whitehead (2008)***), however bootstrapping social network data should only be used carefully as zero edges (which could result from unobserved associations rather than two animals not associating at all) are resampled as zeros across all replications (***Farine and Carter, 2022***). We, therefore, began by assessing whether such algorithms were appropriate for constructing confidence intervals for our chosen global network metrics. In particular, we wished to ensure that the confidence intervals were not too narrow, as this would lead to false positives in comparing networks i.e. finding statistically significant differences where there may be none and therefore having an inflated Type 1 error rate. For example, consider a scenario in which two ecologists independently sample a random subset of individuals from the *same* population of animals. They then construct two social networks and compute network metrics, along with confidence intervals using some statistical methods. If they compare their results and obtain statistically significant results (e.g., via a t-test) and conclude that the two social structures are different, they have made a Type 1 error because, the two samples come from the same population and should therefore not appear statistically significantly different. Any differences are due to the random subsampling of the individuals, but the social structure of the population is the same. We, therefore, wish to confirm that our proposed bootstrapping approach does not exhibit such problematic behaviour (i.e., that analyses based on two subsamples from the same population yield test statistics and p-values consistent with the null hypothesis that the samples arise from populations that have the same social structure as indexed with the various network metrics.) To this end, we began by combining the Bootstrap algorithm with the sub-sampling analysis. See Appendix 1: Assessment of bootstrapping algorithm for results showing correct calibration under the null hypothesis. Then, to examine how uncertainty relates to sample size, we obtained confidence intervals using the bootstrapped samples for each network metric and recorded their width. Finally, we repeated this process ten times and calculated the mean width of the ten confidence intervals against the number of individuals in the network.

### 4. Correlation analysis between node level metrics of partial and full networks

To assess the accuracy of the node level metrics inferred from a given sample, we looked at the correlation between the values of the metrics in the observed sample, and a smaller sub-sample of the empirical data as suggested by ***Silk et al. (2015)***. First, we calculated node-level metrics of degree, strength, betweenness, clustering coefficient, and eigenvector centrality for each node in the observed network. Then we sub-sampled nodes from the observed network at 10%, 30%, 50%, 70%, and 90% levels without replacement and calculated node level metrics for each sub-sample. Finally, we calculated the correlation coefficient between the metric values of the nodes in the observed and partial networks. The process was repeated 10 times at each level of sub-sampling, and the mean and the standard deviation were recorded from the 10 correlation coefficients at each level.

Lastly, we run a regression analysis (***Silk et al., 2015***) to assess how the values of node-level metrics for partial networks relate to their values in the whole network (See Appendix 2: Regression analysis between node level metrics of sub-sampled and observed networks in Appendix).

## Results

### Association Index and Network Formation

Spatial threshold is selected to be 10 meters for mule deer sample and 15 meters for rest of the species samples. The temporal threshold is arbitrarily chosen to be 7 minutes and accounts for delays in signal reception by the GPS devices. For example, if a GPS unit records a location at 09:57 AM, the observations recorded until 10:04 AM will be evaluated for potential interactions. Table 3 shows the values of network summary statistics for each of the five species. The mule deer sample has the highest mean degree and very high mean strength suggesting a dense social network. The elk sample has a maximum diameter with a value of 9 which implies that it will take a maximum of 9 steps to reach from any individual to another in the elk network. The pronghorn sample has the maximum transitivity and mean local clustering coefficient, depicting that any two associates of a pronghorn are likely to be associated to each other. Figure 1 shows the network structures obtained for all five species.

**Figure 1.**
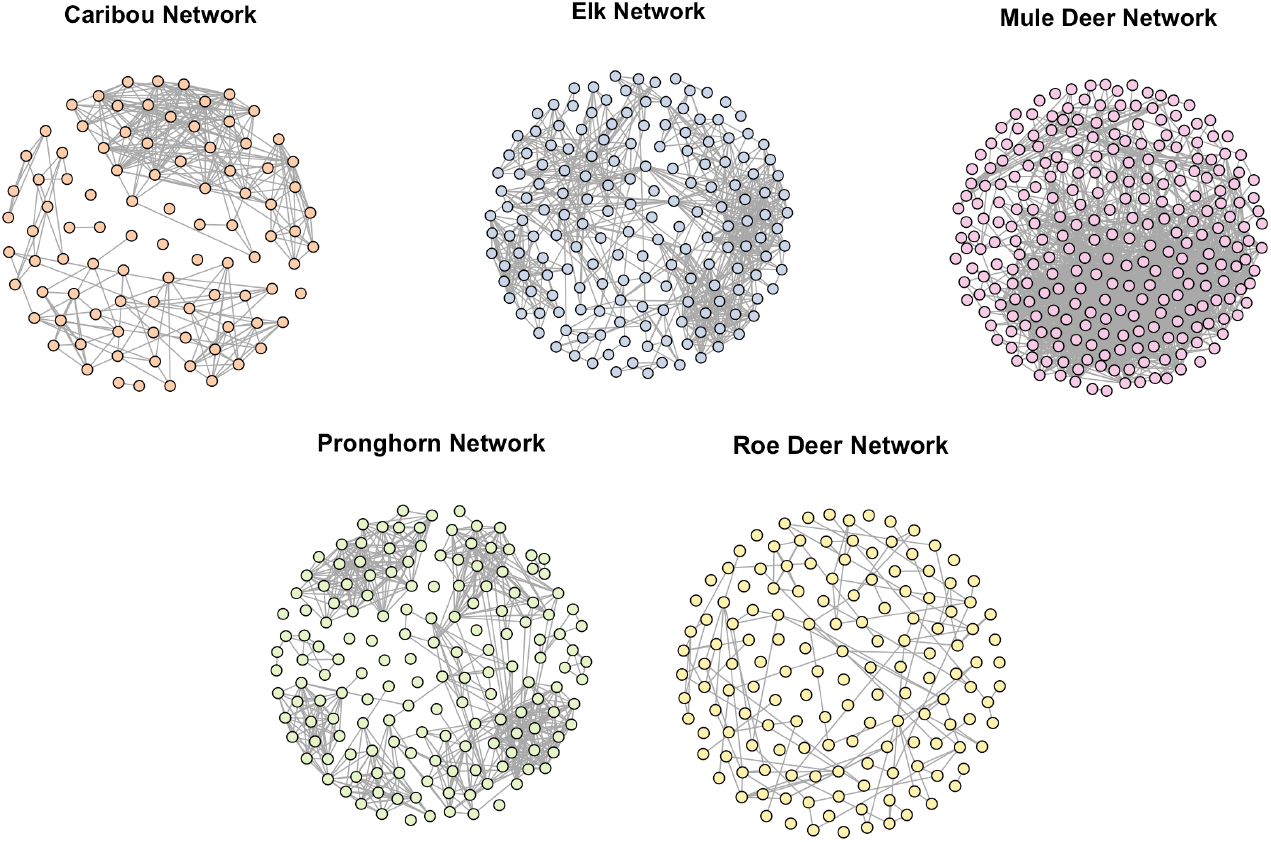
Network structures for the five large herbivores analysed in this study. Each node is an animal, and an edge between two nodes indicates that they have interacted at least once. Note how the pronghorn network is denser than the roe deer network despite having an approximately equal number of nodes. Also, a clear partition in the caribou network can be seen.

**Table 3.**
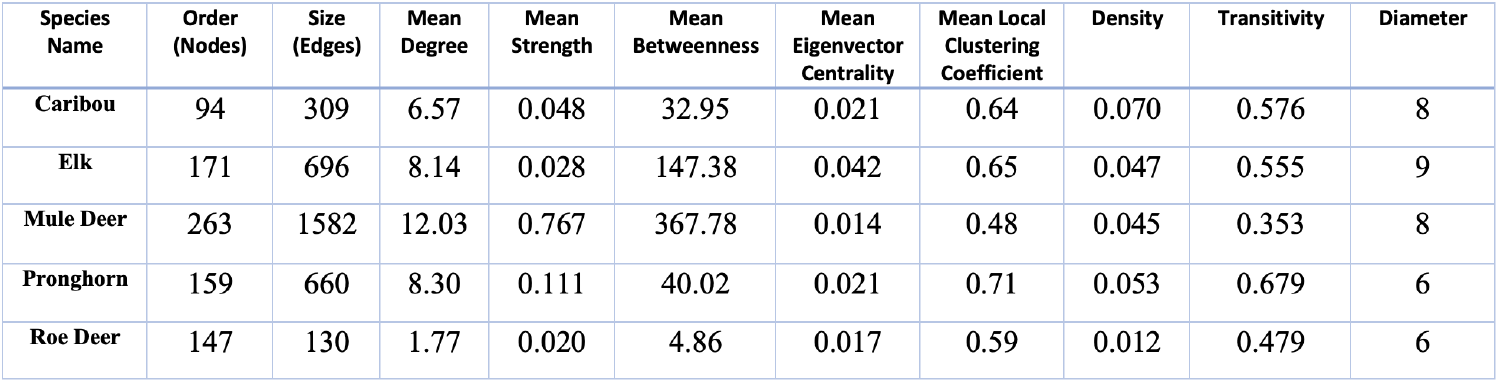
Summary statistics for the networks obtained from the five species. Order of a network represents the number of individuals tagged in the sample with mule deer sample having the greatest order and caribou sample having the smallest order.

| Species Name | Order (Nodes) | Size (Edges) | Mean Degree | Mean Strength | Mean Betweenness | Mean Eigenvector Centrality | Mean Local Clustering Coefficient | Density | Transitivity | Diameter |
| --- | --- | --- | --- | --- | --- | --- | --- | --- | --- | --- |
| Caribou | 94 | 309 | 6.57 | 0.048 | 32.95 | 0.021 | 0.64 | 0.070 | 0.576 | 8 |
| Elk | 171 | 696 | 8.14 | 0.028 | 147.38 | 0.042 | 0.65 | 0.047 | 0.555 | 9 |
| Mule Deer | 263 | 1582 | 12.03 | 0.767 | 367.78 | 0.014 | 0.48 | 0.045 | 0.353 | 8 |
| Pronghorn | 159 | 660 | 8.30 | 0.111 | 40.02 | 0.021 | 0.71 | 0.053 | 0.679 | 6 |
| Roe Deer | 147 | 130 | 1.77 | 0.020 | 4.86 | 0.017 | 0.59 | 0.012 | 0.479 | 6 |

### 1. Pre network data permutations

Pronghorn and roe deer samples capture non-random aspects of the population very well, and for all network metrics, the observed values are significantly different (p < 0.001) from the distribution of permuted network values (Figure 2). Also, for all five species, the mean strength of the observed network are higher than the permuted networks (Figure 2). This indicates that all these samples capture higher association rates than would be expected from a random network under similar assumptions. Researchers dealing with specific study cases and species should not use those network metrics whose observed value lies within the distribution of null values as those do not capture non-random aspects of associations (e.g., transitivity in caribou.)

**Figure 2.**
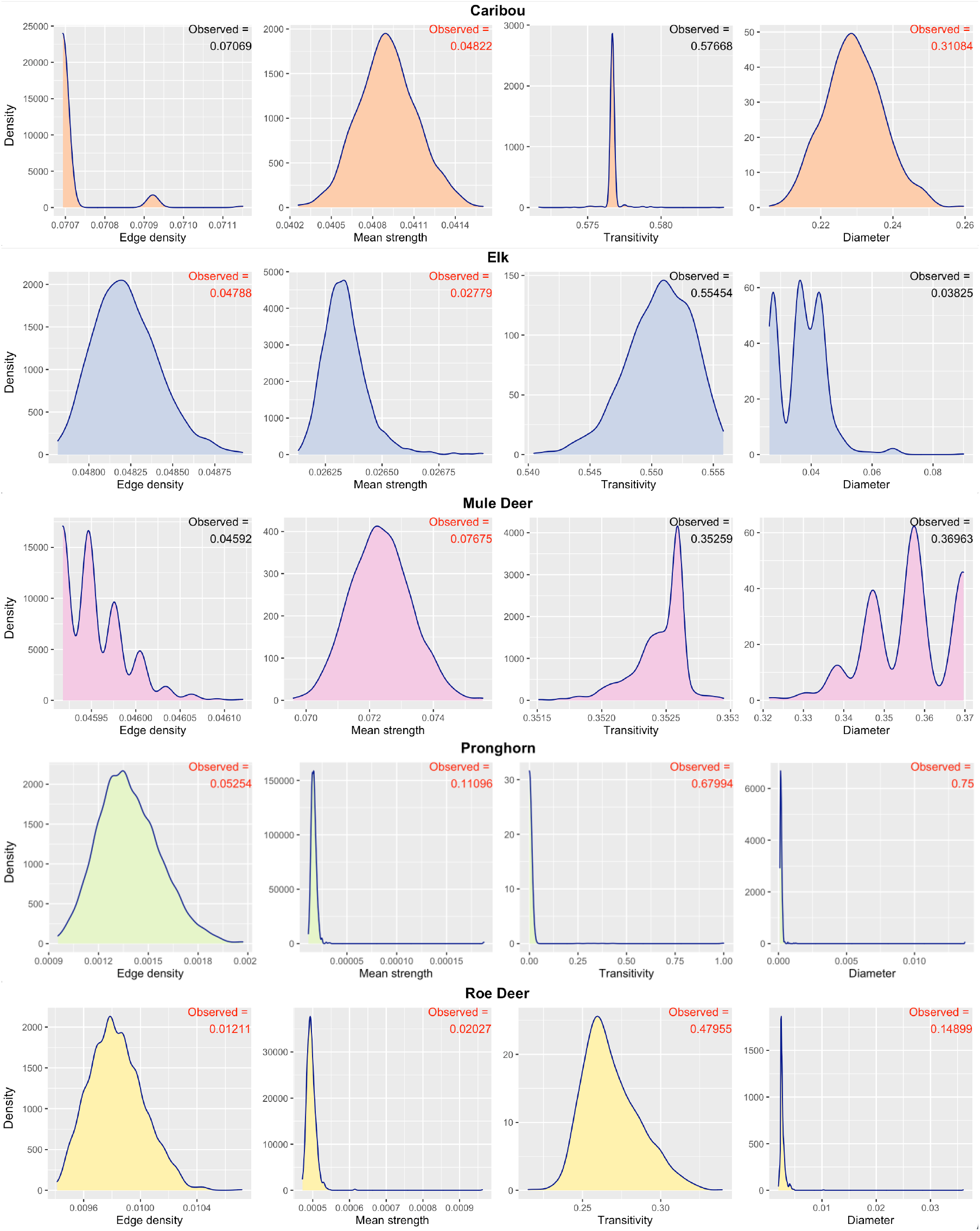
The rows correspond to the five species, while the columns correspond to four standard network metrics. Each plot represents the distribution of network metric values obtained from 1000 permuted versions of the species network. Observed network metric values are quoted at each plot’s top right corner. The color of the observed value (black or red) represents their relative position to the distribution of null values, indicating the extent of non-randomness of the observed metric. The values which are farther from the peak of distribution are displayed in color red and represent that the observed sample captures the non-random aspect well. For example, the observed value of mean strength is high with respect to the distribution of mean strength values from the permuted versions of the data for all five species. This means that all the samples successfully capture non-random interactions between the individuals. The network metrics whose observed value lie within the null distributed values represent that those metric values are not any different from a randomly generated network, such as the density of caribou.

### 2. Subsampling

Performing sub-sampling on the observed samples of networks at various levels revealed the network metrics density and transitivity as most stable (Figure 3.) The uncertainty in these two metrics is comparatively low, even when just 30% of the individuals are present in the sub-sample. Their distribution is centered on the true values, and this allows to estimate bias if the proportion of sampled individuals is known. Transitivity at a sub-sampling level of 10% becomes unreliable for caribou, pronghorn, and roe deer, implying that this metric is a poor measure when the sampling proportion is very small. The bias in mean strength values follows a linear pattern when the subsampled proportion is reduced from 90% to 10% for all five species. The linear pattern suggests that it is possible to correct the bias for mean strength from the sample if the proportion is known. This linear increase must, however, plateau as we approach a census of the population (***Bliss et al., 2014***). The network’s diameter, similar to mean strength, also follows a staircase pattern with lowering sub-sampling levels. However, it does not follow a linear pattern for all five species and tends to plateau. Diameter and mean strength are directly affected by the number of nodes present in the sample. Therefore, care should be taken while using these metrics when the sampling proportion is unknown.

**Figure 3.**
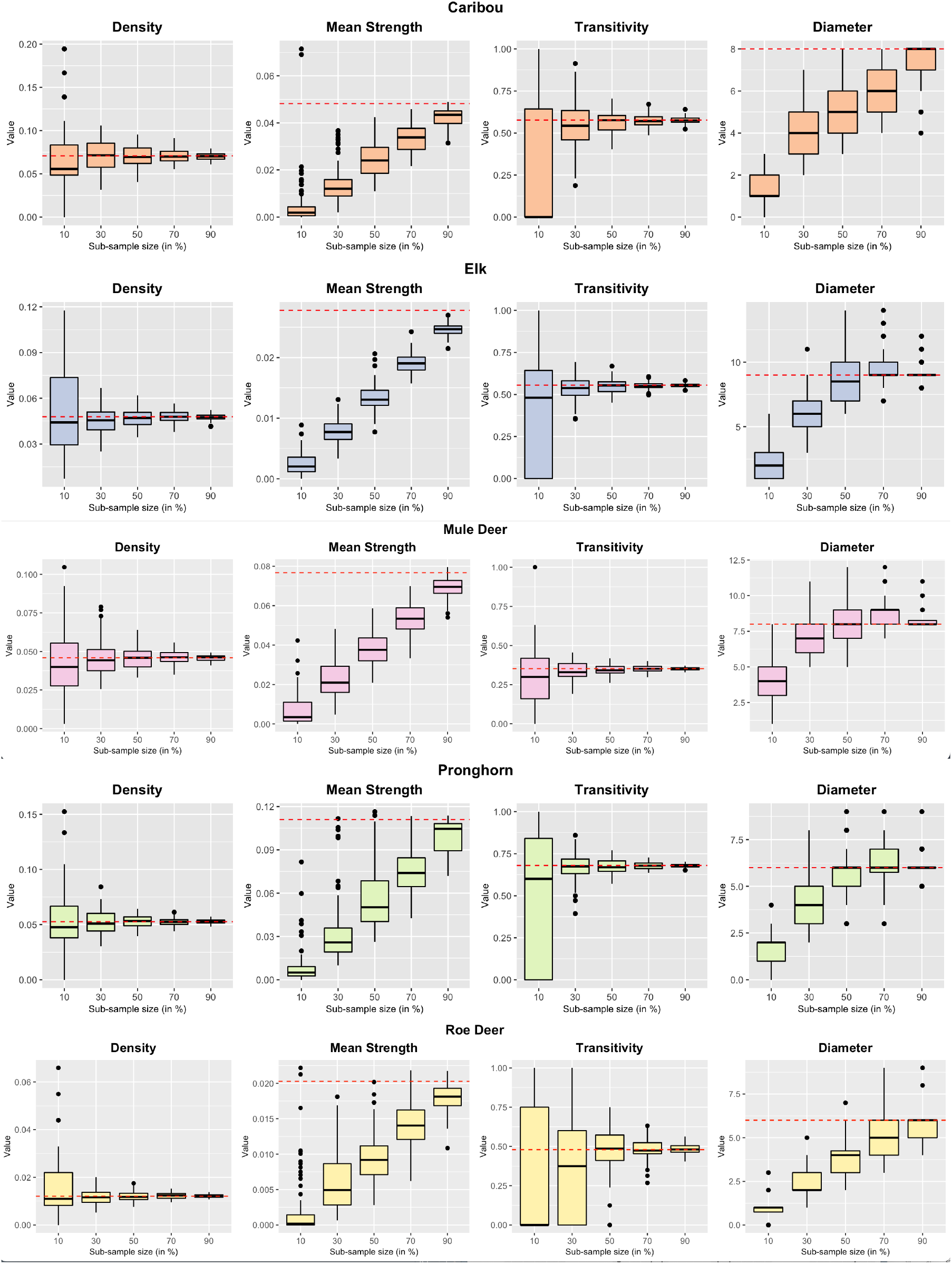
Effect of sub-sampling on four global network metrics. The horizontal red line in each plot represents the metric value in the observed network. The boxplots denote the distribution of network metric values obtained from the observed networks by taking 100 sub-samples at each sampling level. The size of the boxes in the boxplots of a network metric with respect to the sample size represents the extent of uncertainty in that network metric. For example, density has smaller boxes when as low as 10% of the nodes are selected. In contrast, the box size for transitivity is large, representing that density is more stable than transitivity. The extent of deviation of the box position from the horizontal red line depicts the bias in the calculated values of the network metric. The values of mean strength become biased as the sample sizes are lowered. This is because mean strength values are directly affected by the number of nodes present in the network.

We performed subsampling on 1000 permuted versions of the network along with the subsampling on the observed network. The side-by-side visualisation of the network metrics distribution (Figure 4) enabled us to identify the sampling level at which a subsampled network begin to resemble a random network. The plots reveal that for caribou, elk, and mule deer, network metrics density, transitivity, and diameter distribution are identical to that of a null network at all subsampling levels. Nevertheless, mean strength distribution tends to increasingly overlap the distribution of subsamples from the null network when the level of sub-sampling is lower than 90%. For pronghorn, the distribution of all the network metrics obtained from subsamples of the observed network is higher than the values obtained from the subsamples of the null networks at all sampling levels. For roe deer, this is only true for mean strength. The distribution of values for density and transitivity tends to overlap the null distribution at 50% and 30% levels, respectively.

**Figure 4.**
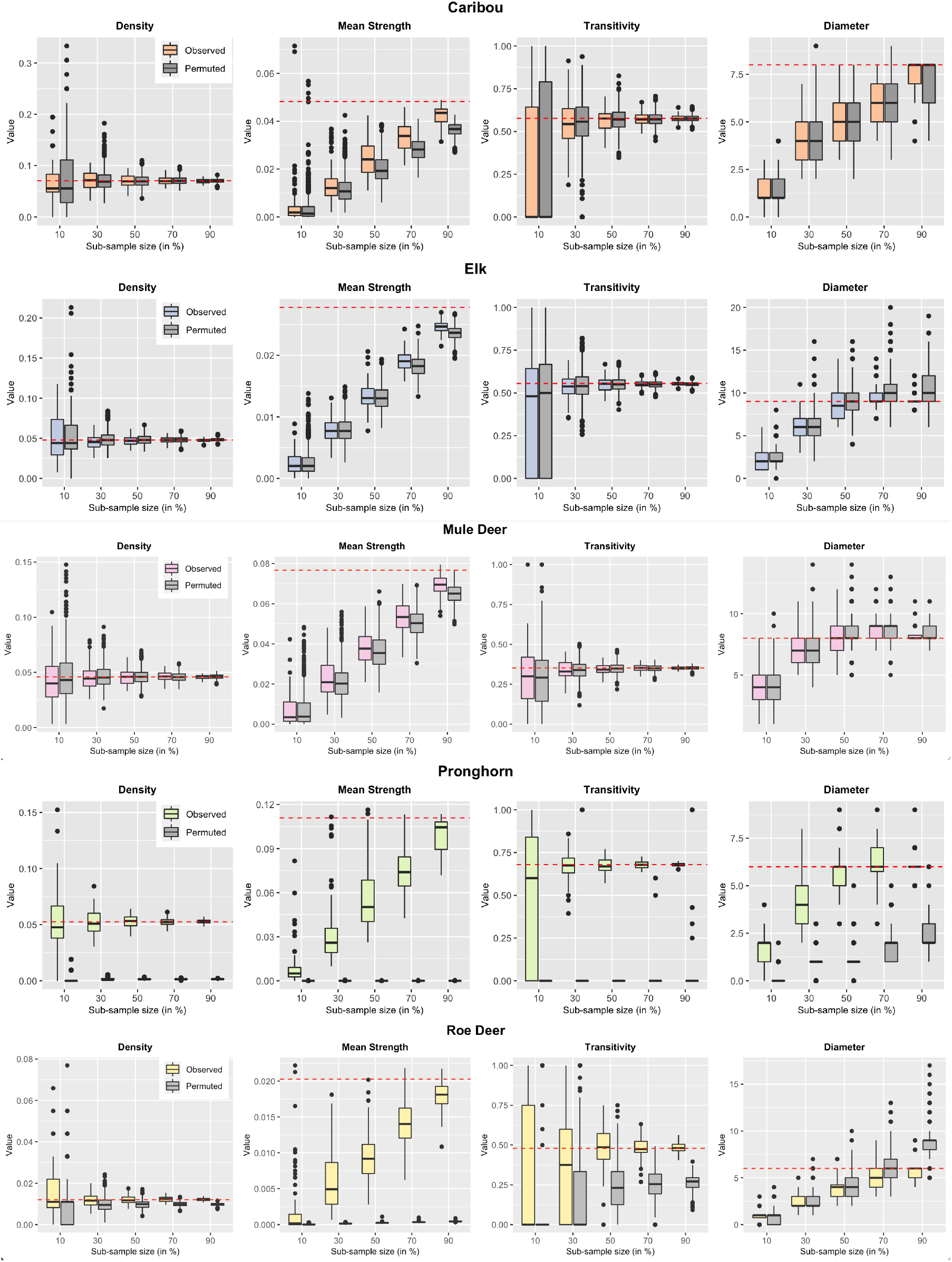
Sub-sampling of permuted networks. The grey boxplots are obtained by calculating network metric values on 1000 permuted versions. The non-grey boxplots are the ones that we obtained in Figure 3. The horizontal red line in each plot represents the observed metric value. Comparing the subsamples of the observed network with those of permuted networks identifies the sample proportion where the non-random aspects of the observed network start looking similar to those of random networks. For example, the mean strength of caribou subsamples in the observed network starts to overlap with the distribution of permuted subsamples at 70% level and becomes almost identical at 10% level. On the other hand, the mean distribution of pronghorn subsamples remains higher than those of permuted subsamples distribution at as low as 10% level.

### 3. Bootstrapping Confidence Intervals

To investigate the extent of uncertainty in the values of network metrics, we used bootstrapping technique to obtain confidence Intervals and plot their width against the sample size (Figure 5 (a)). Confidence intervals can be calculated for any network metric at any sample size. For the network metric density and transitivity, the mean width of the confidence intervals increased with decreasing sample size for all five species. Mean width of density remained comparatively low for as few as 50 samples but began to increase below that value for all five species. For transitivity, the width remained less than 0.2 when at least 100 individuals are tagged for all the species, except for roe deer. The minimum width for roe deer is 0.4, even when all the individuals in the sample are considered. At smaller sample sizes, mean width approaches 1 for all the five species, which is the maximum value transitivity can attain for any network. Therefore, to check that these confidence intervals are not too wide and increase the likelihood of Type 1 errors, we have compared two nonoverlapping sub-samples from the observed sample to check for significant results (See Appendix 1: Assessment of bootstrapping algorithm). We have ensured that the bootstrapping algorithm does not generate spurious statistically significant results.

**Figure 5.**
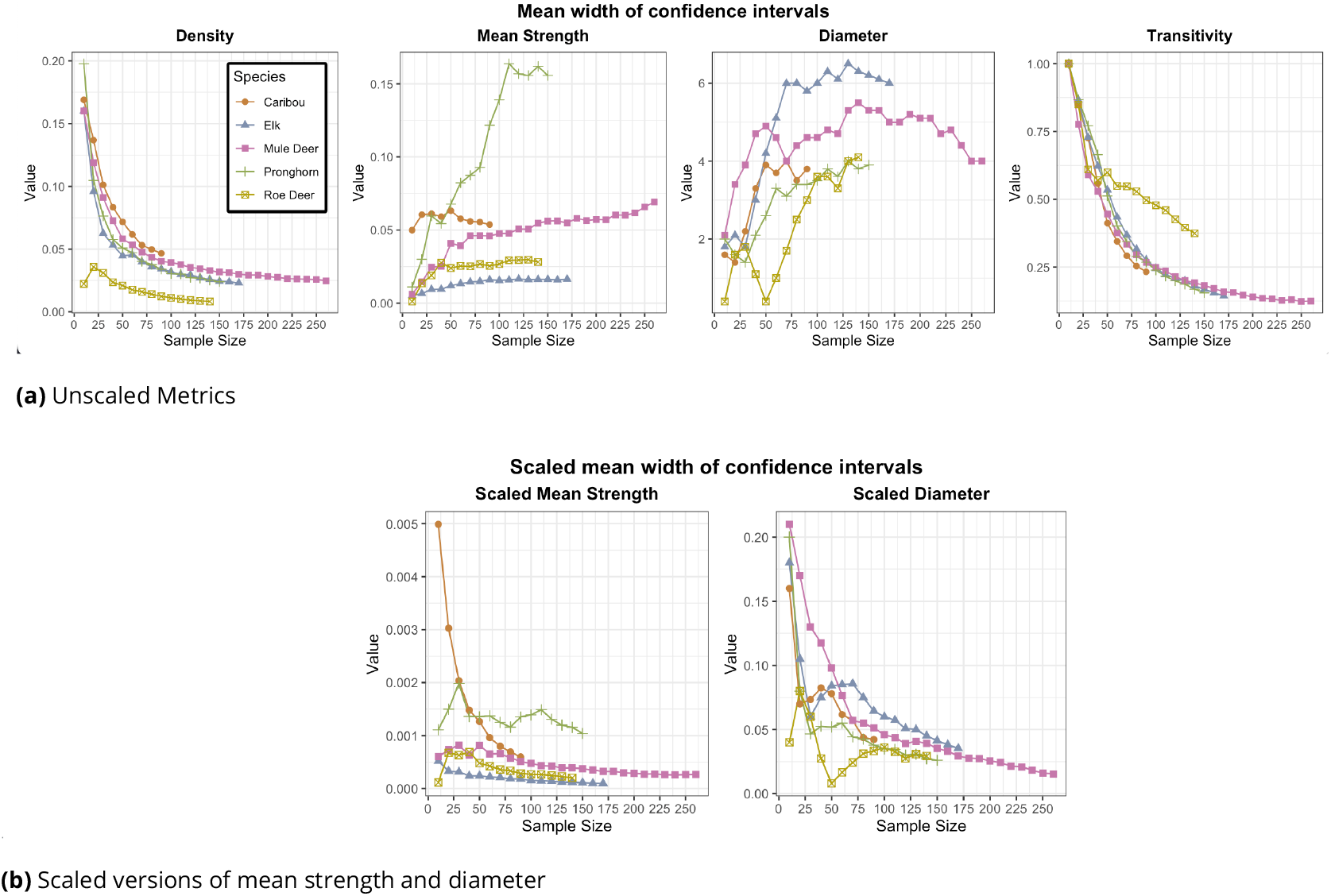
The plots show the mean widths of 95% confidence intervals obtained from bootstrapped sub-samples of a network. The mean widths of density and transitivity increase with lower sample size, which indicates increasing uncertainty around the point estimate of the network metrics. However, the pattern is reversed for mean strength and diameter because the values for these two metrics are directly affected by the number of nodes present in the network. Therefore, we consider scaled versions of these two metrics where the number of nodes at each level scales the values.

For mean strength and diameter, the width of confidence intervals does not increase with a decrease in the sample size. This is because the values of these metrics are directly affected by the number of nodes in the network e.g., say there are N nodes in the network, the possible degrees of a node can be anywhere between 0 and N-1; however, if we remove M (< N) nodes from the network, the new possible value for the degree will lie between 0 and (N-1)-M, which results in a narrower width of the confidence intervals. Therefore it can be helpful to consider the scaled versions of these metrics where they are scaled by the number of nodes in the network (Figure 5 (b)). The scaled versions of these metrics follow a similar pattern to transitivity and density. The width of confidence intervals increases with a decrease in the number of individuals sampled. Some fluctuation in this pattern is observed for the density and scaled diameter for some species, when the sample size is low (e.g., roe deer sample.) Depending on the selection of nodes in the subsample, the value of the diameter in the sub-sampled network can reach extreme values at each level of sub-sampling.

### 4. Correlation Analysis

The correlation of all network metrics between the sub-sampled and observed network declined as the proportion of sub-sampled nodes in the network decreased (Figure 6). However, the pattern and rate of decline are different across network metrics. Degree remains well correlated for mule deer when as few as 10% of nodes are sub-sampled. Mean correlation coefficients of strength, betweenness, and clustering coefficient decline almost linearly with a decrease in the sub-sampling proportion, with slightly more variance in caribou values than mule deer values. For both species, the values for eigenvector centrality became unreliable with high variability even when 90% of the individuals are present in the sub-sample in most of the sub-sampled networks, suggesting that it is a poor measure to use.

**Figure 6.**
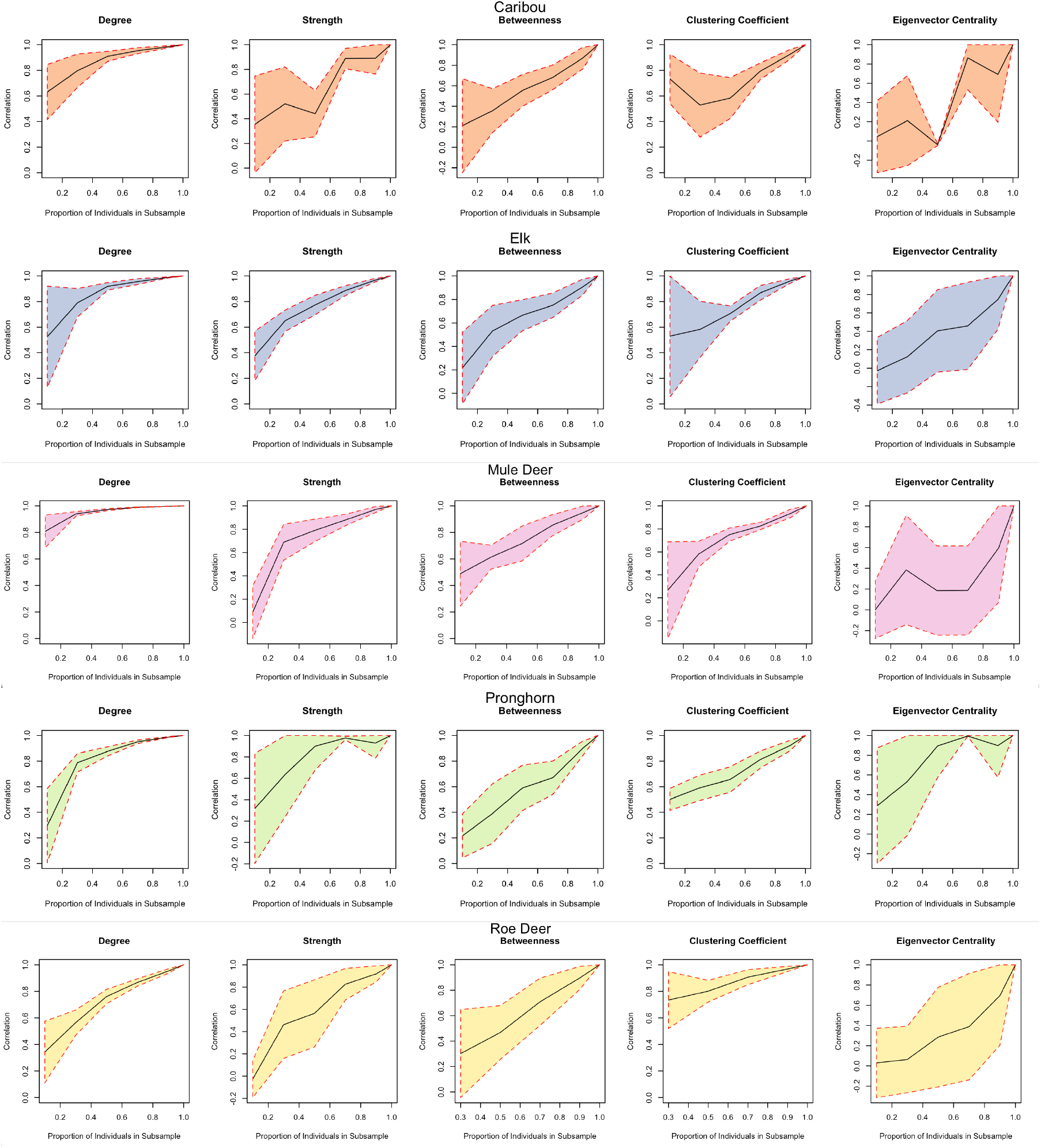
The plots show the correlation of local network metrics between the nodes of sub-sampled and observed networks. The black line in the plots indicates the mean correlation coefficient value between the local metrics of nodes present in the sub-sampled network. The colored region depicts the standard deviation of the correlation values at each sampling level. For example, degree value remains highly correlated with comparatively low standard deviation, even at lower sampling levels.

#### Box 1.

##### Instructions for Ecologists

1. Define the network edges by choosing a sensible distance threshold based on the research question, species sociality, and information obtained from the data (see section Identifying Associations from Raw GPS data for more details.)
2. Check if the interactions captured by the sample are non-random with the help of network permutations. Network metrics can be deemed suitable after assessing whether they capture non-random associations via network permutations.
3. Identify stable network metrics concerning the species and the available sample using sub-sampling from the observed data.
4. Identify the minimum sampling effort required to determine the network properties that are different from a randomly generated network by comparing the sub-sampled networks from permuted data sets with the sub-samples of the observed data.
5. Obtain confidence intervals around the point estimates of network metrics using the bootstrapping algorithm, which also takes care of the autocorrelated structure of telemetry relocation data. The width of confidence intervals can also be analysed for lowering sample sizes.
6. To assess which local network metric remains least affected with lowering sample sizes, obtain a correlation coefficient between the node level metrics from the observed sample and the same nodes from the sub-sample. The local network metrics with a high correlation (>0.7) are expected to be more stable and should be chosen for further analysis as they are more likely to represent the position of individuals in the network, similar to their position in the full population.

## Discussion

Using the four-step paradigm on GPS telemetry observations for five species of ungulates, we assessed the stability of global and local network metrics and obtained measures of uncertainty around the point estimates that can be used to obtain reliable inferences about the structure of a social network. First, using data permutations, we found whether the data collected captured the non-random aspects in all or some of the network metrics: this was entirely true in roe deer, for instance, whereas in other species such as elk the data collected were able to capture non-random associations only with edge density and mean strength. This is a key step in our approach, because at this stage the researcher can make the decisions on whether using such metrics in their study case. Differences among the species in the way metrics responded to permutations analysis most likely reflect different sampling regimes and designs rather than the ecology of the different species. Second, sub-sampling from the observed sample revealed density as the most unbiased measure of animal networks with low uncertainty, even at small sub-sampling proportions. Third, we introduced bootstrapping techniques for animal social networks, which allowed us to compute confidence intervals around the point estimates of the network measures. Density and scaled version of mean strength emerged as two of the most robust network metrics. Fourth and last, correlation analysis between the node level metrics of the observed network and the subsampled network highlighted the network metric degree to be most correlated with the observed network metric values, even at 40% of sub-sampling levels. This means that if the information about sampling proportion is available, the relative degree of each individual can be used to estimate the true degree distribution. Users can take advantage of the functions in the R package, aniSNA (***Kaur, 2023***) to undertake such an analysis of their data. We summarise the steps that should be taken to perform this analysis in Box 1. Instructions for Ecologists.

Despite being a commonly used tool to understand animal ecology (***Hock and Fefferman, 2011***; ***Krause et al., 2010***; ***Robitaille et al., 2019***; ***Silk et al., 2017***; ***Sosa et al., 2021a***,***b***), social network analysis can be challenging when applied to real-life datasets (***Castles et al., 2014***; ***Farine, 2015***). (***Castles et al., 2014***) performed tests to demonstrate a distinction between networks built using different interaction and proximity techniques. Similar tests performed by ***Farine (2015)*** illustrated that the conclusions by ***Castles et al. (2014)*** cannot be generalized across species. A researcher’s choices during the data collection and the analytical stage affect the networks produced. Therefore, the inferences generated may not reflect true characteristics and can be highly sensitive to these decisions (***Ferreira et al., 2020***; ***Castles et al., 2014***). Furthermore, the information available about the sampling protocols may be incomplete or may not be available at all. However, this does not imply that social network analysis should not be conducted on such data. It is prime to use statistical methods that would help extract as much information as possible, along with details about the uncertainties due to partial data and sampling strategies. Performing permutations to randomize autocorrelated GPS data stream (***Farine, 2017***; ***Spiegel et al., 2016***) is a first step to distinguish the best network metrics that capture the non-random aspects of social interactions.

Different network metrics capture different aspects of the network; some networks may have more non-random elements than others, depending on the species’ sociality and the sampling strategies adopted to collect the data. The analyses helped highlighting the network metrics that distinctively capture these non-random aspects. Based on these analyses, we recommend using the network metric mean strength as an assessment metric to identify if the captured interactions are significant enough to generate reliable analysis results. Apart from the four network metrics we chose to work with, it is helpful to run this analysis on multiple network metrics that seem suitable given the research question (e.g., coefficient of the variation of edge weights.) Once a network metric is chosen by the user, further analysis of the available dataset can be carried out to answer the research question. Researchers should keep in mind that our approach is particularly suited (but not limited) to autocorrelated telemetry relocation, although its use could be expanded to more rarefied observational data (e.g., low frequency observations of individually recognizable individuals in a population.)

Caution should be taken while reporting the values of social network metrics when the sample size is small relative to the population (***Lusseau et al., 2008***; ***Farine and Strandburg-Peshkin, 2015***). As a general rule, the smaller the sample size, the more considerable uncertainty can be expected in the observed values. However, some network metrics remain unbiased despite significant uncertainty, whereas others would become biased as the sample size decreases. Sub-sampling from the observed sample and permuted versions of the network revealed helpful information regarding the stability of certain metrics and the proportion of individuals required to ensure a non-null network. For the samples we used, density and transitivity emerged as the more stable metrics, remaining unbiased when as low as 10% and 30% of the individuals were sub-sampled, respectively. Network metrics such as the mean strength became biased as we lowered the number of nodes in the sub-network. However, it was well characterised by a linear relationship which eventually plateaued. The choice of individuals in the sub-sample greatly affected some network metrics such as diameter. However, it always tends to plateau when the proportion of sampled individuals goes up. Sub-sampling from permuted versions of the data and comparing it with the distribution of sub-samples from the observed network revealed the minimum sub-sampling level required to ensure a non-null network structure. ***Davis et al. (2018)*** investigated the effect of sampling effort on the accuracy of social network analysis and concluded that increased sampling intensity may not always increased the accuracy of network measures especially when the sampling regime is already very intense.

Reporting the values of social network metrics point estimates is not enough especially when a large proportion of individuals in the population is not monitored. It becomes equally important to communicate uncertainty around those estimates (***Whitehead, 2008***; ***Lusseau et al., 2008***). We presented bootstrapping as a powerful approach to evaluate confidence intervals around the point estimates of network metrics from the observed data. Bootstrapping enabled us to assess the extent of variation in the network metrics if a different set of individuals was sampled from the population. However, caution should be taken while computing the confidence intervals for those global network metrics, which are directly affected by the network’s number of nodes, such as mean strength. In such cases, the values should be scaled by the number of nodes present in the sample. The network metrics of density and scaled mean strength are those having low uncertainties, even at small sample sizes.

Past studies have tried to uncover how the values of node-level metrics are affected when the sample sizes are lowered (***Silk et al., 2015***; ***Franks et al., 2010***; ***Costenbader and Valente, 2003***). Our work expands on the findings by ***Silk et al. (2015)*** by performing the correlation and regression analyses between the local metric values of the nodes in the entire sample to the values of the same nodes in a smaller sub-sample. With correlation analysis at various subsampling levels, we determined how the correlation rate decreases as the sub-sample size is lowered. Out of the five node level metrics that we tested, degree seemed to work better, with high correlation even at low sample proportions. Also, we recommend not using eigenvector centrality based on a population subsample as the metric being a higher-order statistic lacks robustness and, therefore, is highly sensitive to the selected nodes. This observation agrees with the simulation study by ***Silk et al. (2015)***. We conclude that care should be taken while comparing the metric values between nodes when a small proportion is tagged from the population. Indeed, the social network positions captured by a small sample may not reflect the actual positions in the network in such cases.

The goal of this paper was not to make inferences about the network characteristics of an entire population but to analyse how different network metrics scale under downsampling. As a matter of fact, despite we had access to large telemetry samples from five different species, they represent a subset of an unknown population with unknown size. The methods discussed here can help pinpoint useful social network metrics that remain robust when trying to answer a particular research question. Those metrics that suffer data thinning and become unstable should not be used with telemetry data, which is typically used to monitor a small proportion of the actual population. Also, we used data from multiple species of large herbivores with very different ecology and characteristics, including migratory/ non-migratory and from very social to solitary species. During the analysis, we voluntarily disregarded the ecology of the species, because our goal was not to perform inter-species comparisons and make inferences on their respective social networks but to determine which social network metrics perform well/poorly across the species.

Despite our a priori disregard of the ecology of the five species for the reasons stated above, we found interesting differences among them which deserve to be discussed here. Firstly, the fact that the data collected from a more solitary species such as roe deer (See Table 3) better capture the non-randomness of the association compared to more gregarious species such as elk suggests that sample size (in proportion to the actual population size) should be higher in more gregarious species. In addition, sampling regimes can affect the social network patterns and related ecological inference. For example, a high-density value of the roe deer network as compared to the distribution of null networks could be due to the fact that the sampling was done across six spatially separated capture sites (within 10 x 10 km). This results in very low density values when the data is permuted across these six clusters (Figure 7). Instead, the mule deer’s initial locations (Figure 7) show that the network is already very dense. In the permuted versions of the raw data, the number of random interactions is similar to the number of observed interactions, resulting in observed network density value similar to the density of permuted versions of data. In other words, the sampling strategy (location of the capture sites) may affect the spread of individuals and the density of the respective network structures, therefore researchers need to focus on the ecological interpretation of their social network results after having taken into account of the possible bias introduced by sampling strategies.

**Figure 7.**
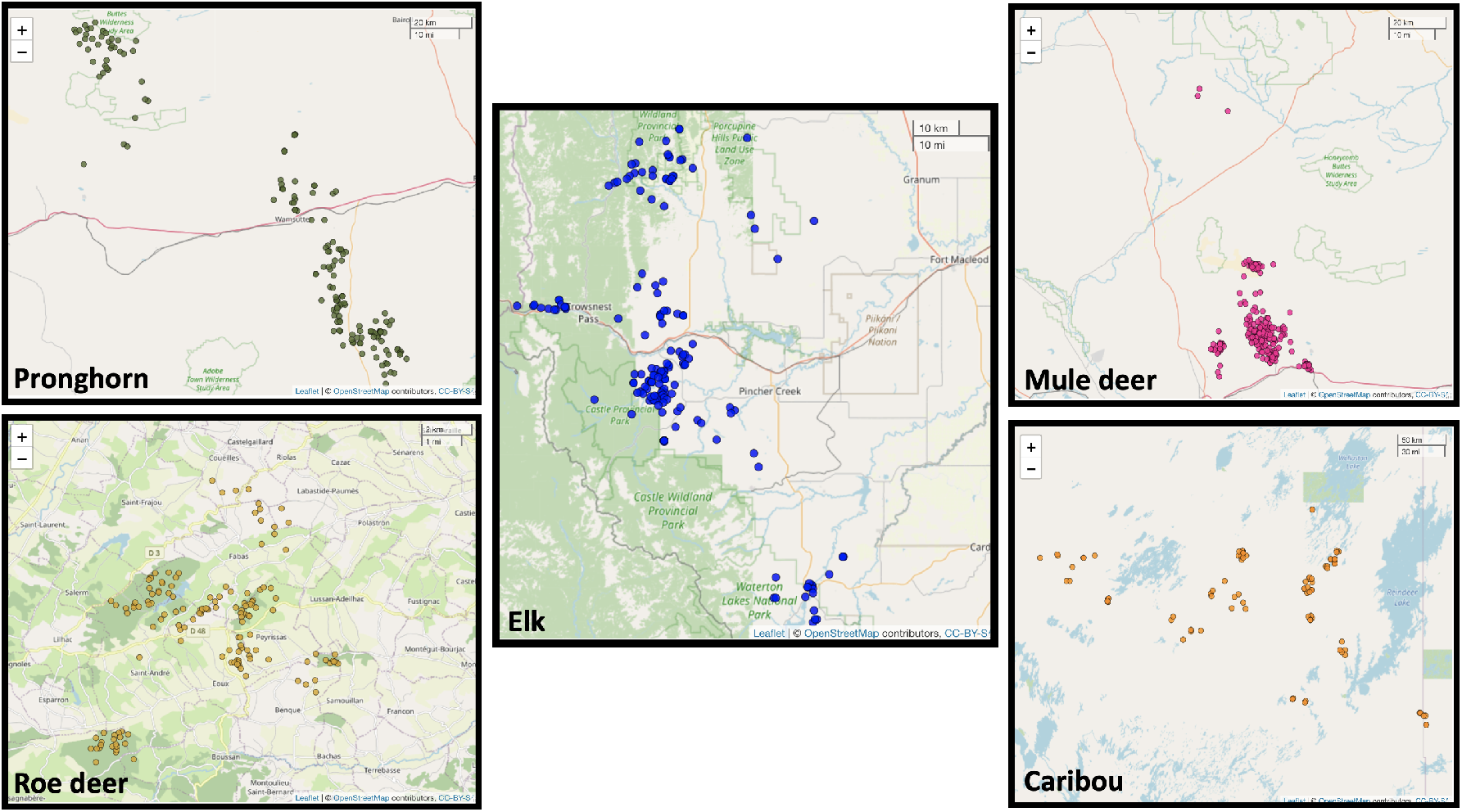
Plot of the initial locations for the individuals belonging to five species in the study. The distribution of these locations explains some of the differences in the values of network metrics. For example, the initial capture locations of mule deer are spatially very close, which is also reflected in the network metric values of the final network of associations. As a result, the mule deer network has the highest mean strength and mean degree (Table 3). In contrast, the roe deer network has the lowest mean strength and mean degree, explained by their six spatially separated capture sites.

Numerous papers have examined the conceptual properties of centrality measures to assist animal social network researchers in selecting the most meaningful and valid measure for their research question and the available data (***Farine and Strandburg-Peshkin, 2015***; ***Franz and Nunn, 2009***; ***Frantz et al., 2009***; ***Borgatti et al., 2006***). The performance of network centrality measures under various sampling regimes and the species sociality could vary to a great extent (***Costenbader and Valente, 2003***; ***Gile and Handcock, 2006***; ***Dawson et al., 2019***) and our work confirms this. Future work involves analysing the effects of observation frequency on the accuracy of network metrics. For example, it could help to understand if it is better to observe individuals for a longer duration with low temporal resolution or a shorter duration with high temporal resolution. In our analysis, sub-sampling on the observed samples is done randomly. However, this aligns differently from the sampling strategies adopted in real life. ***Smith and Morgan (2016)*** investigate the effects on estimates of key network statistics when central nodes are more/less likely to be missing. Application of our methods to determine how the network metrics scale when a different sampling strategy is adopted would be valuable (e.g., whether it is better to sample entire groups, or focus on greater sampling frequency of individuals). Another vital direction forward is to assess the methods presented in this paper to be tested on the GPS telemetry data of the entire population. If data on the whole population is available (e.g., a fenced one), it will be interesting to perform these methods on a subset of that and test if the predictions align with the true values.

Along with all of the advantages to understand animal ecology, SNA presents certain challenges that hinder ecologists from using it to its full extent. We addressed a few of those challenges in this paper and introduced a four-step paradigm to assess the suitability of available data for SNA and extract information for further analysis. The methods are also provided as easy-to-use functions in an R package aniSNA (***Kaur, 2023***). This package allows ecologists to directly apply these statistical techniques and obtain easily interpretable plots to provide statistical evidence for choosing a particular network metric or the choice of individuals tagged for the study. The fact that researchers can compute 95% confidence intervals around their point estimates unleashes new research opportunities, such as tackling specific hypotheses. For instance, researchers can estimate network metrics in a sample population when it is disturbed by human presence to be compared to when it is not disturbed, and the ability to assess the overlap of respective 95% confidence intervals would allow making inference on the effect of human disturbance on sociality. Likewise, this approach can be used to compare social networks within and across populations as a function of temperatures, presence of predators, or different wildlife management strategies, unleashing a range of ecological questions using SNA and related statistical tools.

## Animal welfare

1. Elk in SW Alberta, Canada, were captured (animal care protocol no. 536-1003 AR University of Alberta) during the winters of 2007–2011 using helicopter net-gunning.
2. Pronghorn in southcentral Wyoming, USA, were captured, handled, and monitored according to protocols approved by Wyoming Game and Fish Department (Chapter 33-923 Permit) and University of Wyoming Institutional Animal Care and Use Committee (protocol 20131028JB00037).
3. All animal capture and handling protocols were approved by the Wyoming Game and Fish Department (Chapter 33-937) and an Institutional Animal Care and Use Committee at the University of Wyoming (20131111KM00040, 20151204KM00135, 20170215KM00260, 20200302MK00411).
4. Roe deer in Aurignac, France, were captured (prefectural order from the Toulouse Administrative Authority to capture and monitor wild roe deer and agreement no. A31113001 approved by the Departmental Authority of Population Protection) during the winters of 2005-2012 using drive netting.

## Acknowledgments

1. We thank Damien Farine for thoughtful discussions during the research collaboration at the University of Zurich and providing useful feedback on the manuscript. We also thank Kimberly Conteddu for suggesting improvements on the aniSNA package.
2. This publication has emanated from research conducted with the financial support of Science Foundation Ireland under Grant number 18/CRT/6049. For the purpose of Open Access, the author has applied a CC BY public copyright licence to any Author Accepted Manuscript version arising from this submission.
3. Elk study: We thank the Natural Sciences and Engineering Research Council of Canada (NSERC-CRD), Shell Canada Limited, Alberta Conservation Association (Grant Eligible Conservation Fund), Alberta Sustainable Resource Development, Safari Club International, Alberta Parks and Parks Canada for funding and support.
4. Pronghorn study: We thank Anadarko Petroleum Corporation, Black Diamond Minerals LLC, British Petroleum North America, Devon Energy, Linn Energy, Memorial Resource Development, Samson Resources, Warren Resources Incorporated, the Bureau of Land Management-Rawlins Field Office, Wyoming Game and Fish Department, Wyoming Governor’s Big Game License Coalition, and the University of Wyoming (Department of Ecosystem Science and Management, Office of Academic Affairs, and Wyoming Reclamation and Restoration Center) for funding and support.
5. Any use of trade, firm, or product names is for descriptive purposes only and does not imply endorsement by the U.S. Government.
6. Mule deer study: Funding was provided by the Bureau of Land Management, Hunter Legacy 100 Fund, Knobloch Family Foundation, Muley Fanatic Foundation, National Science Foundation, Pew Charitable Trusts, Safari Club International Foundation, Sitka Ecosystem Grant, Teton Conservation District, The Nature Conservancy, Wyoming Game and Fish Department, Wyoming Governor’s Big Game License Coalition, and the U.S. Geological Survey.
7. Roe deer study: We thank the local hunting associations, the Fédération Départementale des Chasseurs de la Haute-Garonne for allowing us to work in the Comminges, as well as all coworkers and volunteers for help collecting data.
8. Caribou study: We thank the Natural Sciences and Engineering Research Council (Canada) and our industry partners in a Collaborative Research and Development Grant, Environment and Climate Change Canada, Canada Western Economic Diversification, the Saskatchewan Mining Association, and the Government of Saskatchewan.

## Author Contributions

Prabhleen Kaur and Michael Salter-Townshend conceived the ideas and designed the methodology with the help of Simone Ciuti. Prabhleen Kaur analysed the data and wrote the manuscript, edited by Michael Salter-Townshend and Simone Ciuti. Adele K. Reinking, Anna C.Ortega, Anne Loison, Federico Ossi, Francesca Cagnacci, Jeffrey L.Beck, Kamal Atmeh, Mark S. Boyce, Matthew Kauffman, Nicolas Morellet, Philip McLoughlin provided the data and contributed to the revision of the manuscript. All authors contributed critically to the drafts and gave final approval for publication.

## Appendix 1: Assessment of bootstrapping algorithm

We Bootstrapped two different subsamples of a social network at each subsampling percentage. Given that the subsamples were taken from the same population, any significant differences should be attributed to overly confident Bootstrapped intervals and non-significant results were to be expected. We repeatedly sampled pairs of networks at each subsampling level and computed the p-value for a significant difference between the two network metrics, as determined using a two-tailed t-test as per ***Snijders and Borgatti (1999)*** (See 3.Bootstrapped confidence intervals for more details.) If the subsampled networks do indeed provided noisy estimates of the network metric calculated across the population and the Bootstrap algorithm did not return intervals that were too narrow, the B Bootstrapped p-values would not contain more than approximately *α*B values less than *α* (for any significance level *α*). Indeed, the p-values would ideally be uniformly distributed between 0 and 1. We do not perform tests of our bootstrapping approach with regard to power / Type 2 errors because, as with all hypothesis tests, the power to correctly identify real differences in network metrics between two different social networks depends not only on sample size but also on the magnitude of the differences. The larger the sample size and the larger the true difference, the greater the power to identify that the networks differ.

For all the network metrics, p-values for difference between two samples is non-significant for all the five species (Figure 8) which indicates that our bootstrapping approach is not overly sensitive i.e. return too many false positive results when used to compare the network metrics of two networks.

**Appendix 0 Figure 8.**
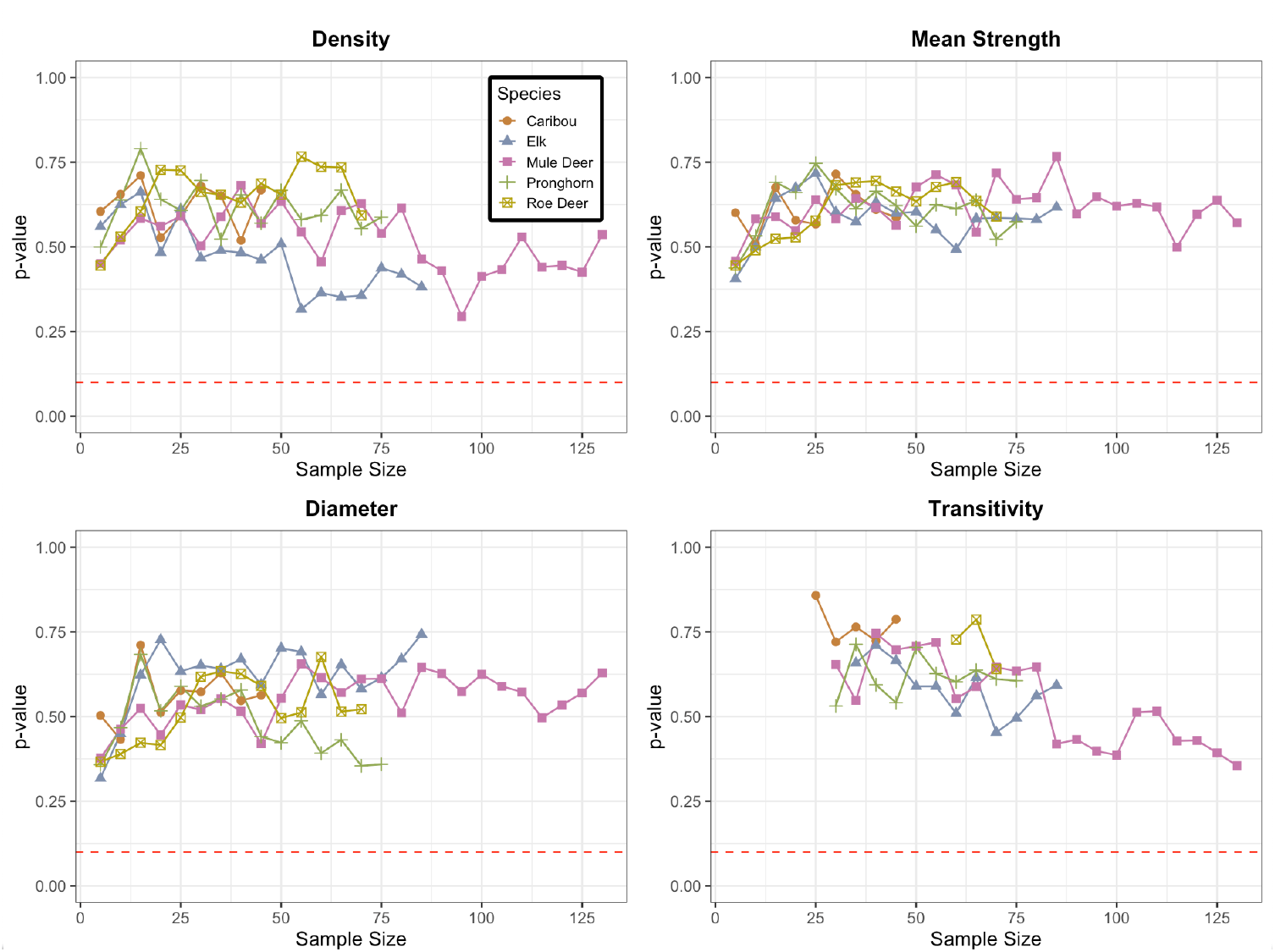
Mean of P Values for the difference in network metrics values for two non overlapping sub-samples of the observed sample. The two non-overlapping sub-samples are bootstrapped 1000 times and p-values are computed for to check for the significance of difference. This process is replicated 10 times to obtain mean of p values. The red dotted line in each plot corresponds to value 0.1 on the y-axis. All the mean p-values lie well above this dotted line.

## Appendix 2: Regression analysis between node level metrics of sub-sampled and observed networks

While analyzing local node level metrics from a partially sampled network, it is useful to know the extent by which the position of an individual in the population controls its position in the sampled network and how this control depends on the choice of individuals in the sample or density of the population.

To answer this, we perform regression analysis such that the values of node-level metrics in the sub-sampled networks are regressed on the values of those nodes in the observed network. As in the correlation analysis, the individuals are further sub-sampled to analyse the effect. Subsampling is done at 10%, 30%, 50%, 70% and 90% levels and is repeated 10 times at each level. The mean slope of regression is calculated for the five network metrics of degree, strength, betweenness, clustering coefficient and eigenvector centrality at each level for each run of the simulation. The slope of regression describes how the value of network metrics calculated in each partial network relates to its value in real network as per ***Silk et al. (2015)***.

The slope of regression is plotted against sampling proportion (Figure 9) The accuracy of node-level metrics from partial networks is highly dependent on the metric being used. In agreement with the results of the simulation study by ***Silk et al. (2015)***, the accuracy of local measures of degree and strength decrease linearly in direct proportion to the proportion of individuals subsampled. In contrast, the accuracy of eigenvector centrality does not depend on that proportion in any of the networks except for elk network. The value of the slope of regression for betweenness decreased non-linearly for four of the species that have low num ber of individuals tagged and followed a near-linear pattern for mule deer that has high number of individuals observed. The accuracy of the clustering coefficient did not decrease till the level of identified individuals was as low as 50% but declined rapidly at different levels after that for each of the species.

***Silk et al. (2015)*** suggests that the strong relationship between the proportion of individuals sampled and the accuracy with which local measures (degree and strength) predict the actual value of an individual’s centrality is notable. This implies that it is possible to correct for this effect if the proportion of sampled individuals in a population is known.

**Appendix 0 Figure 9.**
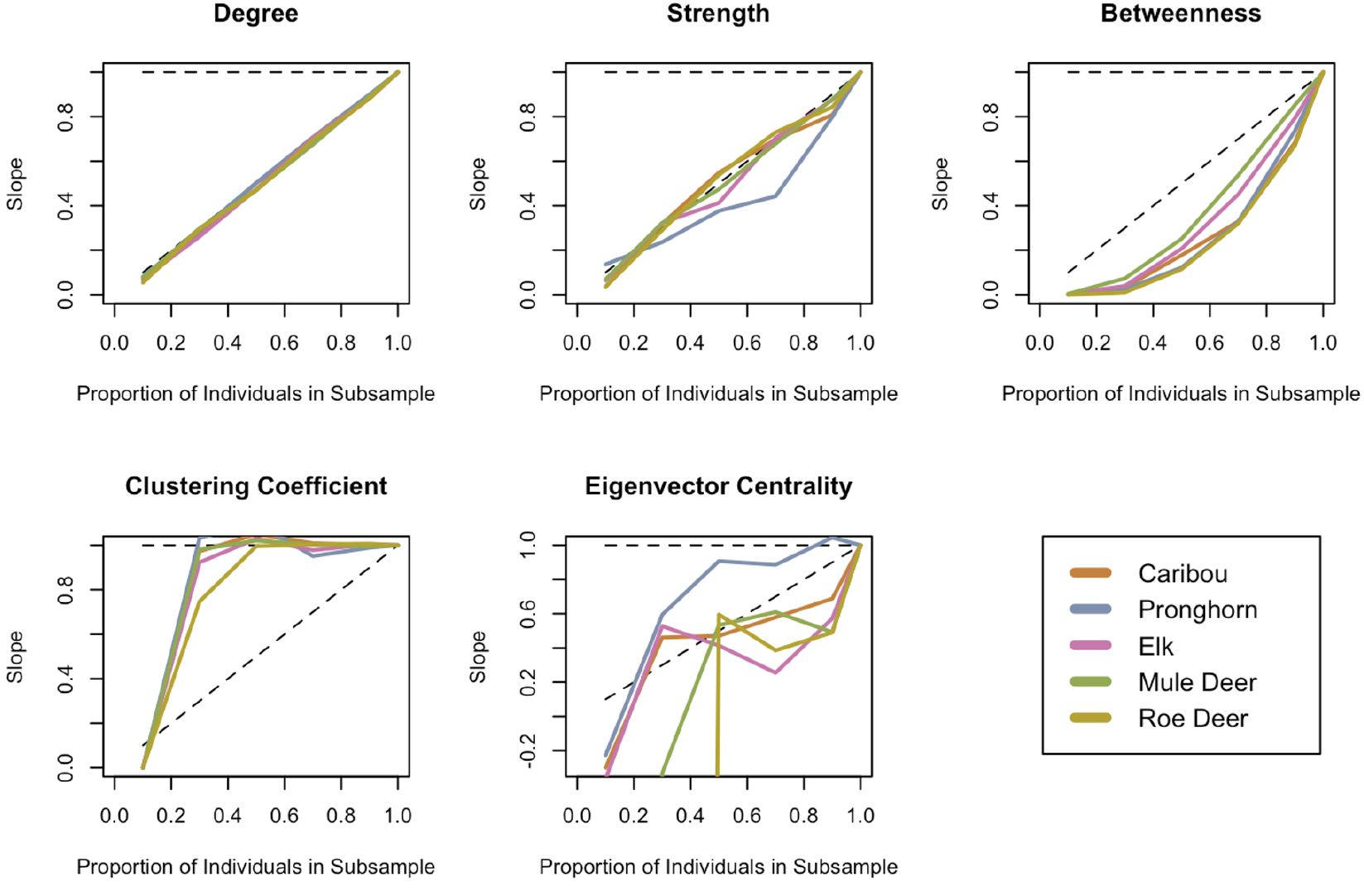
Regression analysis of local network metrics between the nodes of partial and observed network. In each plot, X-axis denotes the proportion of nodes in the sub-sample and Y-axis shows the corresponding value of the slope of regression calculated by regressing the local network metrics of sub-sampled nodes and observed network nodes.

